# Fasting Status and Epigenetic Clock Stability: Implications for Aging Research

**DOI:** 10.64898/2026.06.04.729731

**Authors:** Kirsten Seale, Varun Dwaraka, Ilinca M. Giosan, Tavis Mendez, Ryan Smith

## Abstract

**Background:** Epigenetic clocks are DNA methylation-based biomarkers increasingly used in aging research and clinical trials. A recent assessment of 18 clocks across multiple short-term perturbations concluded that most demonstrate only moderate biological reliability, raising concerns about their translational utility. However, epigenetic clocks differ substantially in their construction and in the biological signals they capture, and their sensitivity to physiological perturbation may not be a flaw but a consequence of the construction. To understand this more clearly, we undertook a focused investigation of biological reliability for a single, well-characterised perturbation, an overnight fast followed by acute refeeding, examining how and why clock estimates may shift with physiological state.

**Methods:** We evaluated 24 epigenetic clocks spanning five construction categories - first and second generation classical clocks (eg. Horvath, Hannum, PhenoAge), the PC versions of the classical clocks, SystemsAge organ-system clocks, mortality-trained clocks (GrimAge, PCGrimAge, OMICmAge), pace of aging clocks (DunedinPACE) and the IntrinClock, across three datasets: a within-person paired fasting design (n = 15 pairs), a cross-sectional cohort of fasted vs non-fasted (n = 2,895), and EPICv2custom technical replicates (n = 96 samples from 4 individuals). For each clock, we quantified the acute fasting effect with and without immune cell adjustment, decomposed between-person and within-person variance at successive adjustment levels (Raw, EAA, IAA), and benchmarked biological variability against the technical measurement floor.

**Results:** Fasting followed by acute refeeding was associated with group-level shifts of 0.5-3 years in immune-sensitive clocks, while within-person reliability remained high (Raw clock ICC median ~0.96). These observations are compatible because fasting effects are small relative to the age-driven between-person variance that dominates the ICC denominator. The magnitude of the observed shift varied by clock. PC transformations showed larger effects than their classical counterparts in the paired cohort (PC Hannum −2.03 vs. Hannum −1.37 years; PC PhenoAge > PhenoAge; PC Horvath > Horvath), SystemsAge showed the largest effects (1.15-2.9 years younger when fasted), and mortality-trained clocks (GrimAge V1/V2, OMICmAge) and DunedinPACE showed no detectable acute effect (all FDR p > 0.10). Immune cell adjustment attenuated or eliminated the fasting effects in sensitive clocks (PC Hannum 88% attenuation; SystemsAge Blood 99.7%); no clock retained a significant fasting effect after FDR-corrected immune adjustment in either cohort. Within the cross-sectional cohort, a clock’s immune content, which is the fraction of its age-independent variance explained by immune cell composition, was correlated with the degree to which immune adjustment attenuated its fasting effect (r = 0.68, p = 0.003). IntrinClock, designed to exclude immune-variable CpGs, showed no fasting effect in either cohort (immune R^2^ = 3.2%), serving as a negative control. Technical replicates confirmed near-perfect measurement reproducibility (median Raw ICC > 0.97), establishing that variance in fasting pairs reflects biology, not noise. Immune-adjusted ICCs behaved differently across clocks in ways consistent with their composition: for clocks where fasting generated within-person variance, immune adjustment removed it and ICC increased (SystemsAge EAA 0.768 to IAA 0.913); for clocks unaffected by fasting, immune adjustment removed between-person structure and ICC fell substantially (OMICmAge 0.922 to 0.160), reflecting the estimation cost of fitting many immune cell predictors to stable residuals. Cross-sectional replication (n = 2,895) confirmed immune cell redistribution at scale. Mortality clocks reached significance cross-sectionally despite resistance to acute fasting.

**Conclusions:** Epigenetic clock responses to an overnight fast followed by acute refeeding varied systematically by training category in our data. PC-based clocks, which concentrate correlated CpG variance including that associated with immune cell composition, showed the largest shifts; mortality-trained clocks showed no detectable acute effect. A framework that summarises a clock by its ICC alone, without identifying the biological source of its within-person variation, can misread structured, perturbation-driven biology as measurement noise. ICC is not a fixed property of a clock, it is shaped by the study design, the population heterogeneity, the perturbation, and the adjustment applied. We recommend that clock reliability be assessed on a perturbation-specific, clock-by-clock basis, with variance decomposition at each adjustment level and explicit benchmarking against technical replicates.

## 1. Introduction

DNA methylation-based epigenetic clocks have emerged as among the most promising biomarkers of biological aging. Since the development of the first pan-tissue clock ^1^ and blood-based clock ^2^, the field has produced increasingly sophisticated algorithms trained on diverse outcomes: mortality ^3^, phenotypic aging ^4^, multi-omic aging ^5^, organ-specific aging ^6^, and the pace of biological aging ^7^. These tools are now used in intervention trials, epidemiological studies, and early clinical applications ^8^, where their value depends on a fundamental property: reliability.

Technical reproducibility of epigenetic clocks is well established. When the same DNA sample is processed repeatedly, intraclass correlation coefficients (ICCs) typically exceed 0.9, indicating minimal technical noise ^9^. Far less attention, however, has been devoted to biological reproducibility, which is the degree to which clock outputs remain stable across repeated sampling from the same individual over short time intervals, during which transient physiological states such as feeding, stress or circadian variation may occur. Sehgal et al. ^9^ evaluated clock stability across meals, stress, and environmental exposures, reporting that most clocks achieve only moderate biological reliability (0.4 - 0.7 for age-adjusted clocks). Based on these findings, the authors concluded this represents a critical limitation to clinical translation, noting that only PC GrimAge consistently achieved ICC values exceeding 0.75.

We argue that this conclusion conflates measurement instability and biological sensitivity. While excellent for evaluating technical reproducibility, reliability metrics alone cannot distinguish between unstructured measurement noise and structured biological responsiveness. The ICC reductions observed in biological replicates could indicate that clocks are imprecise instruments giving noisy readings, or they could indicate that clocks are precise instruments faithfully detecting real physiological change between sampling conditions, and that reduced ICC reflects biological responsiveness rather than instability. Distinguishing these interpretations requires identifying what is driving the between-condition variability.

To distinguish whether observed clock variability reflects an identifiable biological mechanism or is simply unstructured measurement noise, we isolated a single well-characterised perturbation - acute refeeding following an overnight fast. Fasting induces well-characterized changes in circulating immune cell composition ^10,11^. Immune cell composition contributes significantly to epigenetic age acceleration in many widely-used clocks tested to date ^12^. Under this framing, a clock that shifts under acute refeeding following a fast is not unreliable, it is correctly detecting a real, transient change in the biology of the sample.

Crucially, not all clocks should be equally sensitive to immune cell changes. Clocks differ fundamentally in their construction, what features they use, how those features are combined and what outcome they were trained to predict. The first-generation clocks ^1,2^ and the second-generation PhenoAge ^4^ were trained directly on chronological or phenotypic age in cross-sectional cohorts, without explicit handling of cell-composition signal, which means any sensitivity to immune cell flux arises incidentally from their training data rather than by design. The principal component (PC) transformation projects clock CpGs into a lower-dimensional space that captures correlated methylation patterns, including those driven by cell composition ^13^. SystemsAge ^6^, built natively from principal components of organ-specific CpG modules, carries this property by design. By contrast, mortality-trained clocks like GrimAge ^3^ predict lifespan through DNAm surrogates of plasma proteins, and OMICmAge ^5^ integrates CpGs with estimated proteins, metabolites, and clinical measures, both capturing a survival signal that should be largely independent of short-term immune flux. DunedinPACE ^7^ measures the pace of biological aging from longitudinal within-person change, rather than cross-sectional age prediction, and may be less susceptible to acute composition shifts. IntrinClock ^14^, explicitly constructed from CpGs that are consistent across purified immune cell types, should be insensitive to cell composition changes by design.

Given these construction differences, we set out to characterise how acute fasting affects each clock category, examine whether transient immune cell composition changes mediate these shifts, and assess whether each clock’s response could be related to its immune content. We followed three complementary analyses. First, we quantify the group-level effect of acute fasting on clock outputs and immune cell proportions in a within-person paired cohort (n = 15 pairs). Second, we investigate biological reliability by applying an ICC and variance decomposition framework at three adjustment levels (Raw, age-adjusted EAA, immune-adjusted IAA), using technical replicates as a measurement-noise comparator. Third, we replicate the group-level fasting analysis in a cross-sectional cohort (n = 2,895) to investigate the involvement of the immune axis in these shifts.

## 2. Methods

### 2.1 Study Populations

#### Cohort 1: Paired Fasting Analysis (Discovery)

Fifteen healthy adult participants (10 male, 5 female; age range 24–52 years) provided paired blood samples under two conditions within the same day. The first sample was collected in the morning following an overnight fast of at least 12 hours (fasted condition). The second sample was collected 2–4 hours after a standardized meal (non-fasted condition). This within-person design controls for all stable between-person factors (genetics, lifestyle, chronic health), isolating the acute effect of fasting on methylation-derived measures. All samples were preprocessed on the EPICv2custom Illumina array platform.

#### Cohort 2: TruDiagnostic Cross-Sectional (Validation)

The validation cohort comprised 2,895 individuals from the TruDiagnostic clinical database who had self-reported fasting status at the time of blood collection. Of these, 1,445 reported a fasted state and 1,450 reported a non-fasted state. This cohort spans a broad age range (20 - 80 years) with 1,207 females and 1,688 males. This cohort provides substantial statistical power to detect fasting effects, though the cross-sectional design does not permit within-person ICC estimation. All validation samples were processed on the EPICv2custom Illumina array platform.

#### Cohort 3: Technical replicates

To establish a technical measurement noise floor, we analyzed EPICv2custom technical replicates (96 samples from 4 individuals). Because technical replicates contain no biological variability between paired measurements, any within-pair variance represents pure measurement noise, providing an upper-bound on ICC and a lower bound on within-person variance for each clock and each adjustment level. The four technical-replicate donors were profiled as a separate extraction-kit reference panel (Zymo versus Promega) and were not among the 15 paired-cohort participants.

### 2.2 DNA Methylation Profiling

All samples were profiled on Illumina Infinium MethylationEPIC arrays: EPICv2custom (TruDiagnostic). Raw IDAT intensity files were imported using the minfi R package ^15^. Samples were excluded based on quality control using minfi (detection p-value > 0.05) and ENmix QCinfo (percentage of low-quality CpGs and bisulfite intensity); one sample failed ENmix QC and was removed. Beta values were generated using single-sample Noob (ssNoob) normalisation ^16^, which performs background correction and dye-bias correction in a single step without requiring an inter-sample reference. For EPICv2custom samples, replicate probe measurements at the same locus were collapsed to a single value per 10-character locus prefix using the sesame package function *betasCollapseToPfx* ^17^, yielding an EPICv1-compatible probe set, and missing methylation values were imputed using the *imputeBetas* function from the sesame package with the EPIC reference. All three cohorts were processed through this identical pipeline, holding array platform and preprocessing strategy constant across the discovery, validation, and technical replicate analyses.

### 2.3 Epigenetic Clock Computation

We computed 24 epigenetic clocks organized into the following categories:

#### PC-based clocks

PCHorvath, PCHannum, PCPhenoAge, PCGrimAge ^13^. These apply principal component analysis to the CpG sites of classical clocks, projecting them into a lower-dimensional space that captures correlated methylation variation.

#### First and second generation classical clocks

Horvath ^1^, Hannum ^2^, PhenoAge ^4^. The original algorithms from which PC versions were derived.

Organ-system clocks: SystemsAge total and 11 organ-specific clocks (Blood, Brain, Heart, Hormone, Immune, Inflammation, Kidney, Liver, Lung, Metabolic, MusculoSkeletal) from Sehgal et al. ^6^, which are also PC-based.

Mortality clocks: GrimAge V1 ^3^, GrimAge V2 ^18^, OMICmAge ^5^. Pace-of-aging clocks: DunedinPACE ^7^.

Immune-independent: IntrinClock ^14^.

### 2.4 Immune Cell Deconvolution

Immune cell type proportions were estimated from the DNA methylation data using established deconvolution algorithms ^19^. Twelve cell types were estimated: CD4+ naive T cells (CD4Tnv), CD4+ memory T cells (CD4Tmem), CD8+ naive T cells (CD8Tnv), CD8+ memory T cells (CD8Tmem), basophils (Baso), memory B cells (Bmem), naive B cells (Bnv), regulatory T cells (Treg), eosinophils (Eos), natural killer cells (NK), neutrophils (Neu), and monocytes (Mono).

### 2.5 Statistical Analysis

#### Paired Cohort — Fasting Effects

Linear mixed models with a random intercept per participant were fitted at two adjustment levels:

**EAA model:** *Clock* ~ *Fasting_Status*+ *Age*+ *Sex*+ (1|*ParticipantID)*

**IAA model:** *Clock* ~ *Fasting_Status*+ *Age*+ *Sex+*11 *immune cells (1*|*ParticipantID)*

Monocytes were excluded from the immune cell set to avoid collinearity, since cell proportions sum to approximately 1. The IAA model therefore removes both age-related and immune-related structured variation when estimating the fasting effect.

#### Cross-Sectional — Fasting Effects

Linear regression was used in place of mixed models, since the cross-sectional cohort has no repeated measures:

**EAA model:** *Clock* ~ *Fasting_Status*+ *Age*+ *Sex (1*|*ParticipantID)*

**IAA model:** *Clock* ~ *Fasting_Status*+ *Age*+ *Sex+*11 *immune cells (1*|*ParticipantID)*

Note:

DunedinPACE: As a pace-of-aging clock rather than an age predictor, DunedinPACE was modelled without an age term for the paired cohort and the cross-sectional cohort.

IntrinClock: Constructed from immune-invariant CpGs by design, IntrinClock was modelled at the EAA level only and serves as a negative control for the hypothesis that immune adjustment modifies fasting effects.

#### Percent Attenuation

The contribution of immune cell composition to each fasting effect was quantified as:

% *Attenuation* = (1 − β_ *IAA* /β_ *EAA*) * 100

This metric estimates the proportion of the fasting effect explained by shifts in immune cell composition.

#### Immune Content (R^2^)

To quantify the degree to which each clock’s age-independent output depends on immune cell composition, we regressed EAA on all 11 immune cell proportions in the cross-sectional cohort (n = 2,895). For this analysis, EAA residuals were computed as:

*Clock* ~ *Age* + *Sex*

to remove sex-linked immune differences that could confound the clock-immune relationship (sex differences in immune cell proportions are well-established and would inflate the immune R^2^ of clocks with sex-associated CpGs). The resulting R^2^ represents the fraction of each clock’s age- and sex-residualized variance explained by immune cell composition. We also computed Pearson correlations between each clock’s EAA and each individual immune cell type to identify which cells drive the relationship. For DunedinPACE, age was not included in the model.

#### Multiple Testing Correction

All p-values within each analysis (clock effects, immune cell effects) were adjusted using the Benjamini-Hochberg false discovery rate (FDR) procedure. FDR correction was applied separately within the primary clock set (PC, SystemsAge, Mortality, Pace; n = 20) and the secondary classical-clock set (Horvath, Hannum, PhenoAge; n = 3) to avoid the larger primary set dominating the correction for the smaller classical set. IntrinClock was treated as a negative control and not included in either FDR set; its raw p-value reported uncorrected.

#### Test-Retest Reliability (ICC)

Intraclass correlation coefficients were computed using the psych R package, selecting the ICC2 variant (two-way random effects, single measures, absolute agreement) for the paired and technical replicate cohorts ^20^, using the following formula:

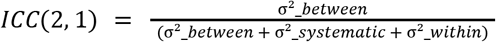

Where σ^2^_between is the between-participant variance, σ^2^_systematic is the variance attributable to systematic differences between the two measurement conditions (fasted vs non-fasted), and σ^2^_within is residual within-person variance. Higher values indicate greater within-person stability relative to between-person spread.

For each clock, ICC was calculated at three adjustment levels to isolate different sources of variation:

- Level 1 — Raw Clock: direct methylation-based clock output, reflecting all sources of variation including chronological age, immune cell composition, and residual biological and technical variation.
- Level 2 — EAA (Epigenetic Age Acceleration): residuals from a regression of Clock ~ Age, removing between-person differences attributable to chronological age and isolating age-independent biological variation.
- Level 3 — IAA (Immune-Adjusted Acceleration): residuals from a regression of Clock ~ Age + 11 immune cells, removing both age- and immune-composition-driven between-person variance.

Rationale for ICC interpretation: Each successive adjustment removes structured between-person variance from the total variance denominator. Because ICC equals between-person variance divided by total variance, removing structured between-person variance necessarily reduces ICC, even if within-person variance remains unchanged.

For the ICC and variance decomposition analyses, EAA residualisation uses Age only. Sex is a stable between-person characteristic that does not vary within individuals across fasting states; including it in the residualisation would remove between-person variance unrelated to measurement reliability, mechanically deflating ICC without adding information about clock stability. Sex is instead included as a covariate in the fasting effect models above, where it improves estimation precision.

#### Variance Decomposition

For each adjustment level, between-person variance (BP) was computed as the variance of per-person means across the two fasting states, and within-person variance (WP) as the mean of per-person variances. These map directly onto the ICC numerator and denominator so the same three adjustment levels (Raw, EAA, IAA) produce parallel BP and WP trajectories that reveal which variance component each adjustment removes.

### 2.6 Software

All analyses were conducted in R (version 4.3+) using lme4 and lmerTest for mixed models, irr and psych for ICC computation, and tidyverse for data manipulation and visualization.

## 3. Results

Study design and cohort demographics are summarised in Figure 1. We used three cohorts: a within-person paired fasting design (n = 15) for primary effect estimation and ICC analysis, a cross-sectional validation cohort (n = 2,895) for replication and immune content quantification, and EPICv2custom technical replicates (n = 96 samples from 4 individuals) to establish the measurement noise floor. The results are organised around four observations: i) fasting produced group-level shifts in epigenetic age that varied by clock training category, were attenuated by immune cell adjustment, and replicated in a cross-sectional cohort, ii) these directional shifts coexisted with high within-person ICCs because individual shifts are small relative to the large age-driven between-person variance, and iii) the PC transformation amplified the fasting effect of every matched first or second generation classical clock, and the Raw-to-EAA ICC drop tracked the fasting effect size at the category level; and iv) direct assessment of the estimated immune cell proportions confirmed fasting-induced cell redistribution as the likely mediator, with each clock’s immune content correlating with the degree of attenuation under immune adjustment.

**Figure 1.**
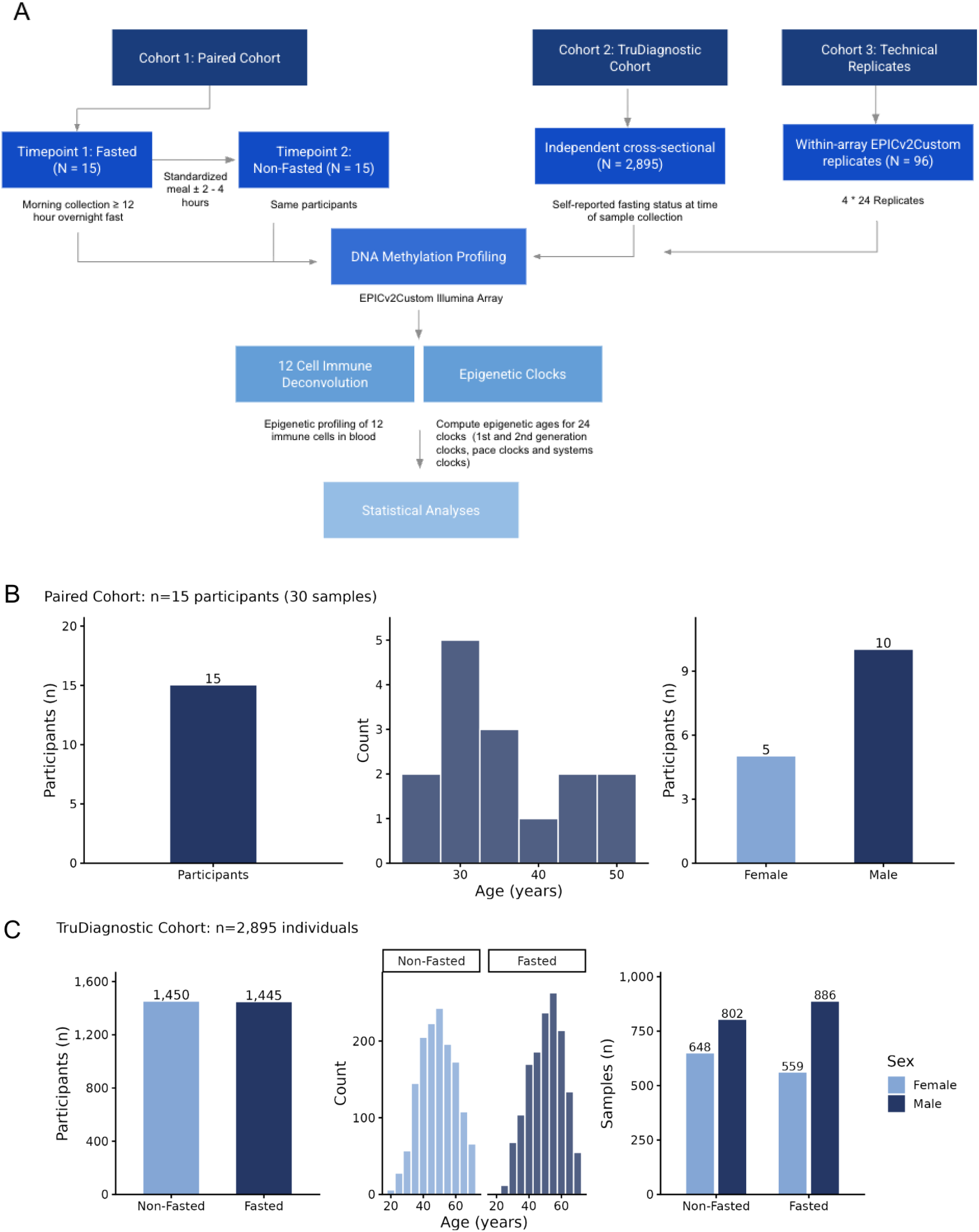
Study Design and Cohort Demographics. Schematic overview of the study design. A: Schematic overview of the study design including three cohorts from left to right: Paired cohort, TruDiagnostic Cross-sectional Validation Cohort, TruDiagnostic Technical Replicates. B: Paired cohort (n = 15): participant counts, age distribution, and sex composition. C: TruD validation cohort (n = 2,895): equivalent demographic summary.

### 3.1 Fasting produces structured group-level shifts that vary by clock training category and are attenuated by immune adjustment

In the paired cohort (n = 15), PC-based clocks exhibited the largest and most consistent shifts when fasted (Figure 2A, Table 1). PC Hannum shifted −2.03 years (FDR p = 0.012) and PC Horvath shifted −1.22 years (FDR p = 0.012). SystemsAge Total, built natively from principal components of organ-specific CpG modules, showed the largest effects among aggregate clocks (β = −1.61, FDR p = 0.038), and 8 of 11 organ sub-clocks reached significance after FDR correction (β across significant sub-clocks ranging from −1.30 to −2.90 years). Mortality-trained clocks, in sharp contrast, were resistant to fasting effects. GrimAge V1 (β = +0.14, FDR p = 0.720) and GrimAge V2 (β = −0.07, FDR p = 0.886) showed no detectable effect, as well as OMICmAge, which displayed a non-significant trend in the opposite direction (β = +0.63, FDR p = 0.110). DunedinPACE was similarly unaffected (β = −0.012, FDR p = 0.530). These clocks predict mortality through DNAm surrogates of plasma proteins (GrimAge) or multi-omic features (OMICmAge) whose between-person variation reflects chronic health status rather than acute immune redistribution. IntrinClock, which is immune-independent by design, showed no fasting association in the paired cohort (β = +0.67, p = 0.364).

**Table 1.** Fasting Effects on Epigenetic Clocks — Paired Cohort (N = 15)

| Clock | EAA $\beta$<br>(years) | EAA SE | EAA p | EAA p<br>(FDR) | IAA $\beta$<br>(years) | IAA SE | IAA p | IAA<br>(FDR) | %<br>Attenuation |
| --- | --- | --- | --- | --- | --- | --- | --- | --- | --- |
| PC Horvath | -1.223 | 0.286 | <0.001* | 0.012* | -0.149 | 0.401 | 0.719 | 0.972 | 87.8% |
| PC Hannum | -2.027 | 0.520 | 0.002* | 0.012* | -0.241 | 0.382 | 0.547 | 0.972 | 88.1% |
| PC PhenoAge | -1.667 | 0.799 | 0.056 | 0.074 | -0.229 | 0.833 | 0.788 | 0.972 | 86.3% |
| Horvath | 0.207 | 0.708 | 0.775 | 0.775 | 1.877 | 1.391 | 0.201 | 0.604 | -808.0% |
| Hannum | -1.368 | 0.564 | 0.029* | 0.088 | -0.517 | 0.759 | 0.513 | 0.757 | 62.2% |
| PhenoAge | -1.185 | 1.128 | 0.311 | 0.467 | 0.625 | 1.977 | 0.757 | 0.757 | 47.3% |
| PC GrimAge | -0.652 | 0.362 | 0.093 | 0.110 | 0.326 | 0.405 | 0.440 | 0.972 | 49.9% |
| GrimAge V1 | 0.136 | 0.327 | 0.684 | 0.720 | 0.670 | 0.609 | 0.293 | 0.972 | -393.9% |
| GrimAge V2 | -0.069 | 0.472 | 0.886 | 0.886 | 0.871 | 0.809 | 0.303 | 0.972 | -1164.1% |
| OMICmAge | 0.626 | 0.343 | 0.089 | 0.110 | 1.168 | 0.615 | 0.084 | 0.972 | -86.5% |
| SystemsAge (Total) | -1.610 | 0.619 | 0.021* | 0.038* | 0.414 | 0.533 | 0.454 | 0.972 | 74.3% |
| SystemsAge: Blood | -2.085 | 0.546 | 0.002* | 0.012* | 0.007 | 0.530 | 0.990 | 0.990 | 99.7% |
| SystemsAge: Brain | -2.896 | 0.787 | 0.002* | 0.012* | -0.060 | 0.606 | 0.923 | 0.972 | 97.9% |
| SystemsAge: Inflammation | -2.034 | 0.721 | 0.014* | 0.034* | 0.063 | 0.546 | 0.911 | 0.972 | 96.9% |
| SystemsAge: Heart | -1.314 | 0.585 | 0.041* | 0.064 | 0.552 | 0.558 | 0.344 | 0.972 | 58.0% |
| SystemsAge: Hormone | -1.299 | 0.438 | 0.010* | 0.029* | 0.122 | 0.567 | 0.832 | 0.972 | 90.6% |
| SystemsAge: Immune | -1.440 | 0.654 | 0.045* | 0.064 | -0.146 | 0.753 | 0.850 | 0.972 | 89.9% |
| SystemsAge: Kidney | -2.080 | 0.773 | 0.018* | 0.038* | -0.059 | 0.456 | 0.899 | 0.972 | 97.1% |
| SystemsAge: Liver | -2.617 | 0.813 | 0.006* | 0.025* | 0.164 | 0.481 | 0.742 | 0.972 | 93.7% |
| SystemsAge: Metabolic | -2.369 | 0.763 | 0.008* | 0.026* | -0.394 | 0.461 | 0.416 | 0.972 | 83.4% |
| SystemsAge: Lung | -1.664 | 0.639 | 0.021* | 0.038* | -0.144 | 0.755 | 0.852 | 0.972 | 91.3% |
| SystemsAge: MusculoSkeletal | -1.150 | 0.493 | 0.035* | 0.059 | -0.224 | 0.408 | 0.596 | 0.972 | 80.5% |
| DunedinPACE | -0.012 | 0.016 | 0.477 | 0.530 | -0.024 | 0.032 | 0.464 | 0.972 | -101.0% |
| IntrinClock | 0.669 | 0.714 | 0.364 | - | - | - | - | - | - |

Note: $\beta$ = fasting effect in years (negative = fasted appears epigenetically “younger”). EAA model: Clock ~ Fasting + Age + Sex + (1|PID). IAA model: additionally adjusted for 11 immune cell types. % Attenuation = $(1 - |\beta_{IAA}|/|\beta_{EAA}|) \times 100$ . Significant effects are denoted with an \* and values are bold.

**Figure 2.**
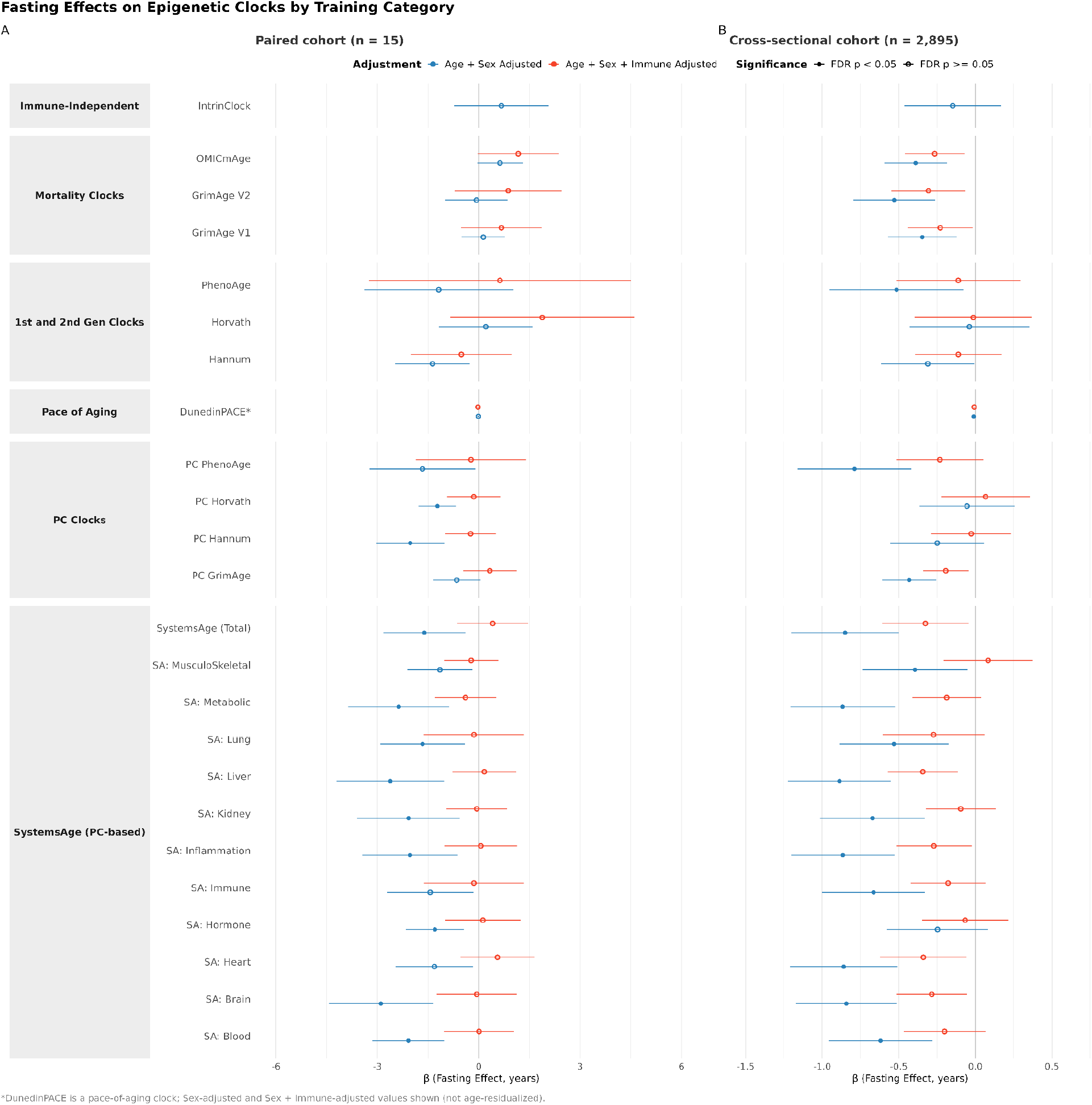
Effect of fasting on epigenetic clock outputs with and without immune adjustment — paired cohort (left) and cross-sectional cohort (right). Side-by-side forest plots of fasting effects (beta coefficients, years) on epigenetic clocks in each of the cohorts, organised by clock family. Blue = EAA model (clock output residualised for age and sex); Red = IAA model (additionally residualised for immune cell composition). Filled symbols indicate p < 0.05; open symbols indicate p ≥ 0.05. Error bars = 95% confidence intervals. Reference group: Non-Fasted. Negative beta = epigenetically younger when fasted. Linear mixed-effects model with random intercept per participant for the paired cohort only.

Adding immune cell proportions to the model (IAA) substantially reduced or eliminated the FDR-significant fasting effects. PC Hannum attenuated by 88% (β: −2.03 to −0.24), PC Horvath by 88% (β: −1.22 to −0.15), SystemsAge Total by 74%, and the significant SystemsAge organ sub-clocks by 83% to 99.7%. After FDR correction, no clock retained a significant fasting effect at the IAA level in the paired cohort (all IAA FDR p > 0.6). The clock-by-clock breakdown of attenuation and its relationship to each clock’s immune content are explored in detail in section 3.4 of the results.

To test whether the sensitivity gradient generalised beyond 15 participants, we analysed the TruDiagnostic cross-sectional cohort with self-reported fasting status. The PC > 1st and 2nd generation classical > mortality > immune-independent gradient held in this cohort. PC PhenoAge shifted −0.79 years (FDR p < 0.001), SystemsAge Total shifted −0.85 years (FDR p < 0.001), and 10 of 11 SystemsAge organ sub-clocks reached FDR significance at EAA. PC Horvath and PC Hannum did not reach significance at this sample size, consistent with their smaller absolute effects. Mortality clocks, resistant to fasting in the paired design, reached significance at population scale (GrimAge V1 −0.35, V2 −0.53, OMICmAge −0.39 years; all FDR p ≤ 0.004; DunedinPACE −0.011 units, FDR p = 0.001), and IntrinClock again showed no fasting association (β = −0.15, FDR p = 0.40), paralleling the paired cohort.

Interpretation of this cross-sectional mortality-clock signal is not straightforward, since the cross-sectional cohort utilises a self-reported fasting metric at a single time point that does not specify duration or pattern, people who report being fasted may differ systematically from those who do not in ways unrelated to fasting itself, and no cross-sectional fasting effect survived FDR-corrected immune adjustment (all IAA FDR p ≥ 0.07). We unpack these caveats in the Discussion.

### 3.2 Excellent within-person reliability between biological replicates coexists with group-level shifts

Despite these group-level shifts, Raw ICCs in the paired cohort remained uniformly high (median 0.96, range 0.93 - 0.98, all clocks except DunedinPACE at 0.81; Table 3, Figure 3A; Supplementary Figure S1), comparable to technical replicates processed through the same pipeline (median Raw ICC > 0.97; Supplementary Table S1). EAA ICCs declined to a heterogeneous good-to-excellent range (median 0.79), with OMICmAge scoring the highest EAA ICC of 0.922. Six clocks (SA: Brain, SA: Liver, SA: Lung, SA: Inflammation, SA: Metabolic, IntrinClock) fall into the 0.64–0.75 range, flagged as below the threshold for “good” reliability ^9^. Immune adjustment shifted ICCs in opposing directions across clocks (Table 3, Figure 3 B,C). For five clocks, ICC rose under IAA, including SystemsAge Total (EAA ICC: 0.768 to IAA ICC: 0.913), and four of its sub-clocks, including SA: Heart, SA: Liver, SA: Metabolic, and SA: Brain (Supplementary Figure S1). The remaining 17 clock ICCs declined from EAA to IAA to varying degrees. The largest declines were SA: Immune (EAA ICC: 0.768 to IAA ICC: 0.000) and OMICmAge (EAA ICC: 0.922 to IAA ICC: 0.160), as well as Horvath, Hannum, and PhenoAge. The PC versions of these classical clocks declined less sharply, for example PC PhenoAge ICC declined from 0.858 to 0.451, compared to Horvath which declined from 0.834 to 0.173. PC Hannum had the largest paired-cohort fasting effect (β = −2.03) but its IAA ICC still fell. PC GrimAge was the only clock whose IAA ICC was essentially unchanged from EAA (0.791 to 0.754).

**Table 2.** Fasting Effects on Epigenetic Clocks — TruDiagnostic Validation Cohort.

| Clock | EAA $\beta$ (years) | EAA SE | EAA p | EAA (FDR) | IAA $\beta$ (years) | IAA SE | IAA p | IAA (FDR) | % Attenuation |
| --- | --- | --- | --- | --- | --- | --- | --- | --- | --- |
| PC Horvath | -0.054 | 0.159 | 0.733 | 0.766 | 0.066 | 0.148 | 0.654 | 0.716 | -22.2% |
| PC Hannum | -0.249 | 0.156 | 0.111 | 0.135 | -0.027 | 0.133 | 0.839 | 0.878 | 89.1% |
| PC PhenoAge | -0.788 | 0.19 | **<0.001\*** | **<0.001\*** | -0.232 | 0.145 | 0.110 | 0.195 | 70.5% |
| Horvath | -0.039 | 0.199 | 0.846 | 0.846 | -0.014 | 0.195 | 0.944 | 0.944 | 64.7% |
| Hannum | -0.311 | 0.155 | **0.045\*** | 0.058 | -0.111 | 0.145 | 0.443 | 0.599 | 64.3% |
| PhenoAge | -0.514 | 0.222 | **0.021\*** | **0.030\*** | -0.111 | 0.207 | 0.591 | 0.715 | 78.4% |
| PC GrimAge | -0.432 | 0.09 | **<0.001\*** | **<0.001\*** | -0.192 | 0.076 | **0.011\*** | 0.072 | 55.5% |
| GrimAge V1 | -0.346 | 0.114 | **0.002\*** | **0.004\*** | -0.229 | 0.108 | **0.035\*** | 0.079 | 33.7% |
| GrimAge V2 | -0.530 | 0.136 | **<0.001\*** | **<0.001\*** | -0.306 | 0.124 | **0.013\*** | 0.072 | 42.2% |
| OMICmAge | -0.390 | 0.104 | **<0.001\*** | **<0.001\*** | -0.265 | 0.099 | **0.008** | 0.072 | 32.0% |
| SystemsAge (Total) | -0.85 | 0.18 | **<0.001\*** | **<0.001\*** | -0.326 | 0.144 | **0.023\*** | 0.077 | 61.6% |
| SystemsAge: Blood | -0.619 | 0.173 | **<0.001\*** | **<0.001\*** | -0.2 | 0.136 | 0.142 | 0.233 | 67.6% |
| SystemsAge: Brain | -0.842 | 0.167 | **<0.001\*** | **<0.001\*** | -0.285 | 0.118 | **0.016\*** | 0.072 | 66.2% |
| SystemsAge: Inflammation | -0.864 | 0.172 | **<0.001\*** | **<0.001\*** | -0.27 | 0.126 | **0.032\*** | 0.079 | 68.7% |
| SystemsAge: Heart | -0.859 | 0.179 | **<0.001\*** | **<0.001\*** | -0.339 | 0.144 | **0.019\*** | 0.072 | 60.6% |
| SystemsAge: Hormone | -0.247 | 0.168 | 0.142 | 0.164 | -0.066 | 0.143 | 0.642 | 0.716 | 73.1% |
| SystemsAge: Immune | -0.664 | 0.171 | **<0.001\*** | **<0.001\*** | -0.178 | 0.124 | **0.152** | **0.233** | 73.2% |
| SystemsAge: Kidney | -0.672 | 0.174 | **<0.001\*** | **<0.001\*** | -0.096 | 0.116 | 0.411 | 0.590 | 85.8% |
| SystemsAge: Liver | -0.887 | 0.171 | **<0.001\*** | **<0.001\*** | -0.342 | 0.117 | **0.003\*** | 0.072 | 61.4% |
| SystemsAge: Metabolic | -0.866 | 0.174 | **<0.001\*** | **<0.001\*** | -0.186 | 0.114 | **0.103** | **0.195** | 78.5% |
| SystemsAge: Lung | -0.532 | 0.182 | **0.004\*** | **0.005\*** | -0.272 | 0.17 | **0.109** | **0.195** | 48.8% |
| SystemsAge: Musculoskeletal | -0.394 | 0.175 | **0.025\*** | **0.033\*** | 0.083 | 0.148 | 0.574 | 0.715 | 78.9% |
| DunedinPACE | -0.011 | 0.004 | **0.001\*** | **0.001\*** | -0.007 | 0.003 | **0.028** | 0.079 | 34.1% |
| IntrinClock | -0.147 | 0.162 | 0.362 | 0.397 | - | - | - | - | - |

Note: $\beta$ = fasting effect in years (negative = fasted appears epigenetically “younger”). EAA model: $\text{Clock} \sim \text{Fasting} + \text{Age} + \text{Sex}$ . IAA model: additionally adjusted for 11 immune cell types. % Attenuation = $(1 - |\beta_{\text{IAA}}|/|\beta_{\text{EAA}}|) \times 100$ . Significant effects are denoted with an \* and values are bold.

**Table 3.**
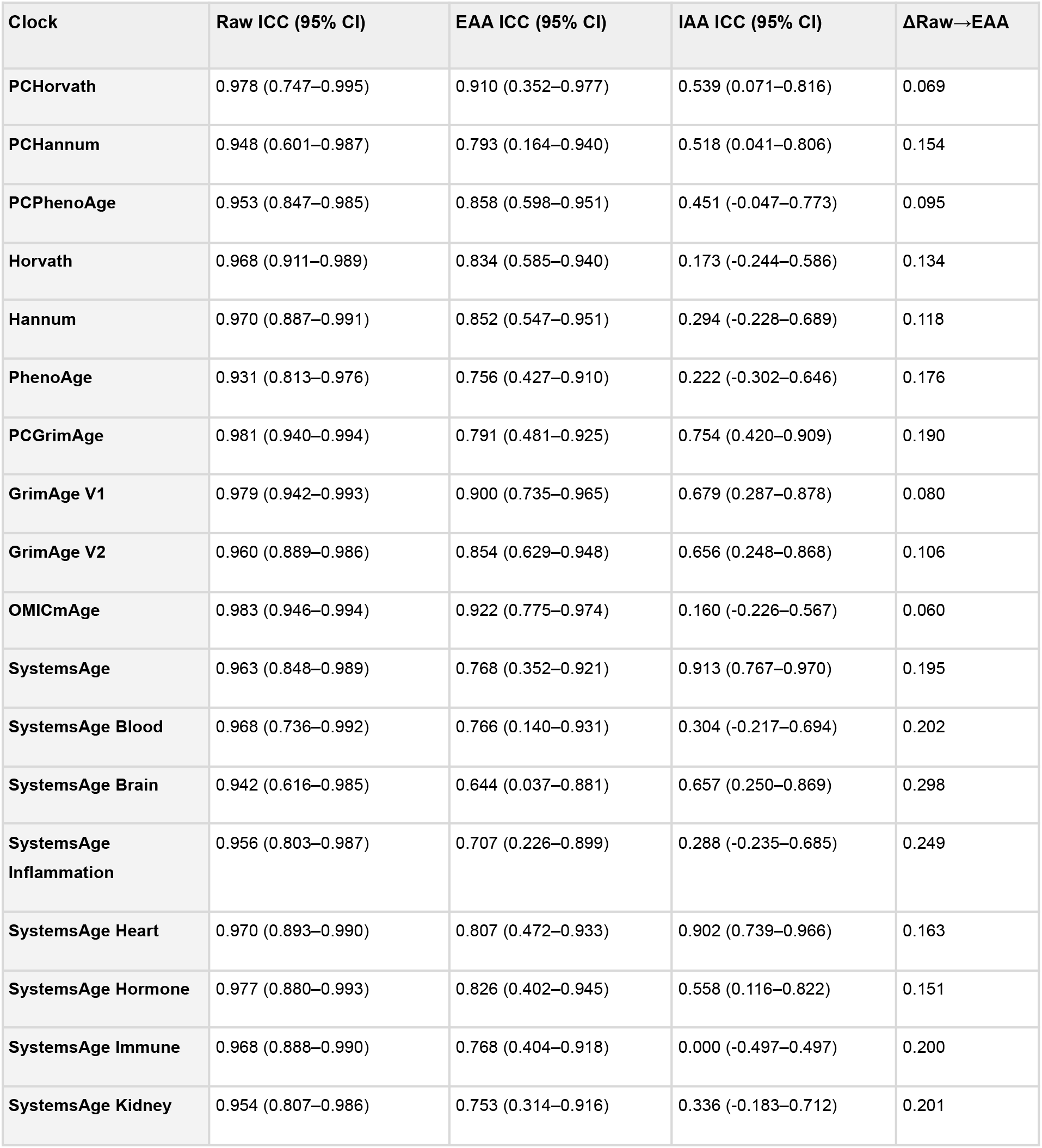
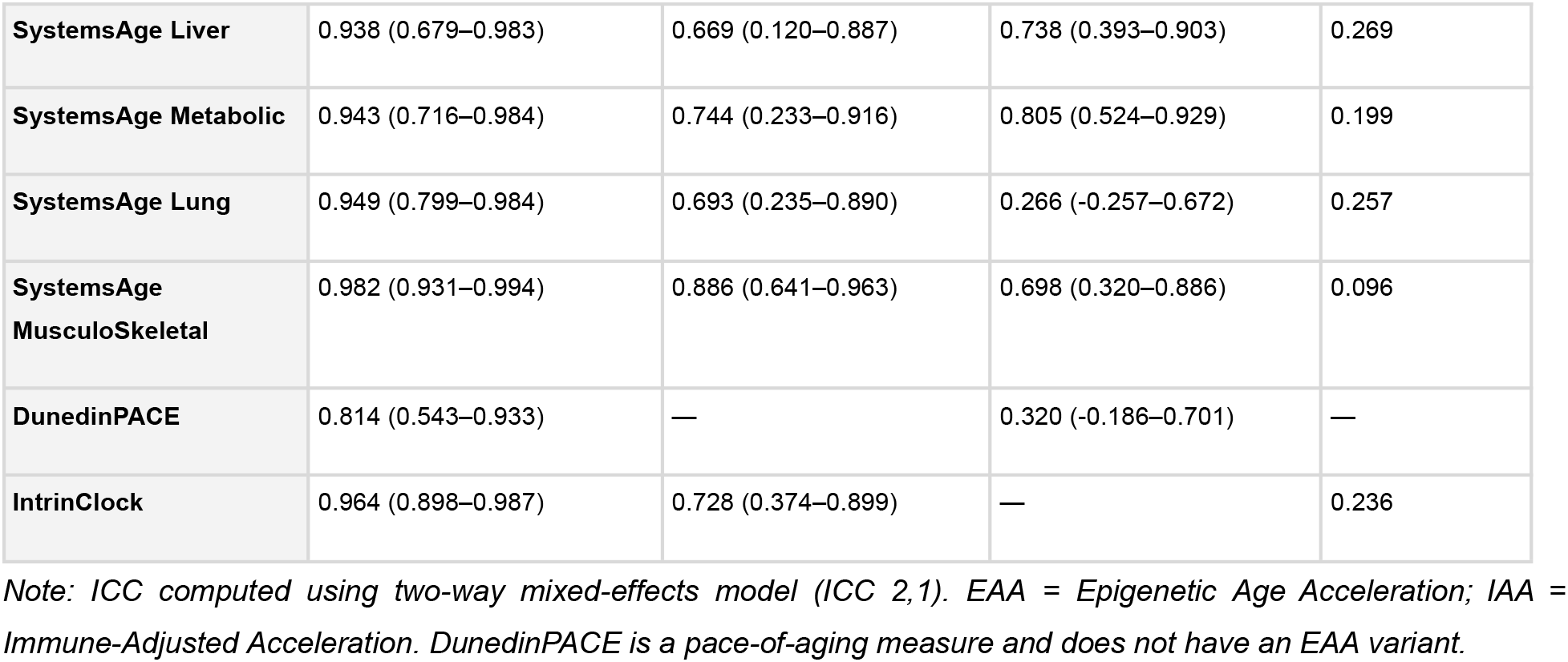
Intraclass Correlation Coefficients (ICC) Across Adjustment Levels — Paired Cohort (N = 15)

**Figure 3.**
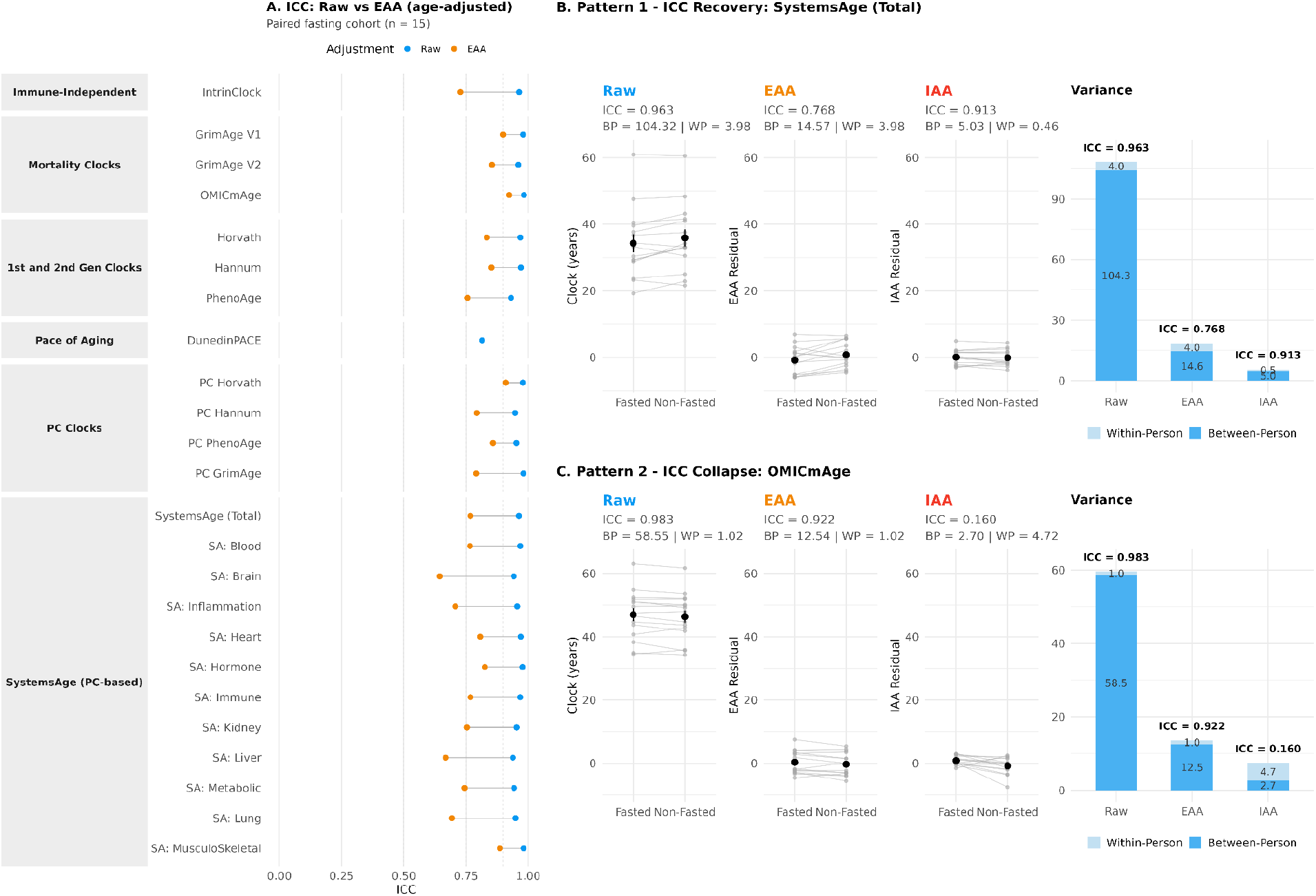
Test-Retest Reliability and Variance Decomposition Across Adjustment Levels - Paired Cohort. (A) ICC (point estimates) for all 24 clocks at two adjustment levels (Raw = blue, EAA = orange), grouped by training category. Dashed vertical lines: ICC = 0.5, 0.75, 0.9. Segments connect Raw to EAA for each clock. The IAA level is omitted here because it is not a clock-reliability metric (see Supplementary Figure S1 for the full Raw + EAA + IAA version). (B) Pattern 1 - ICC recovery (SystemsAge Total): raw and EAA trajectories show a clear fasting-driven shift; at IAA the shift is removed and ICC recovers (0.768 to 0.913) as within-person variance falls. Between-person (BP) and within-person (WP) variance are shown in the adjacent stacked bar. (C) Pattern 2 - ICC collapse (OMICmAge): no visible fasting shift at any level; immune adjustment collapses BP variance from 12.54 to 2.70 (78%) while WP rises from 1.02 to 4.72 from overfitting 11 immune predictors to stable residuals, so ICC falls from 0.922 to 0.160 despite no change in biology. Together the two patterns illustrate that ICC change under immune adjustment is a diagnostic of clock composition.

SystemsAge Total and OMICmAge serve as contrasting examples of how ICC responds to successive variance adjustment. In each case, the resulting ICC change depends on (a) how between-person (BP) and within-person (WP) variance shift at each adjustment level (Raw to EAA to IAA), and (b) whether the BP and WP components themselves are altered by the fasting perturbation (Figure 3B, C). Mechanistic interpretation of these ICC patterns is in section 4.3 of the Discussion.

### 3.3 PC transformations amplify clocks’ sensitivity to fasting

We used the paired cohort to directly compare the matched PC and first and second generation classical versions of the same clocks (Figure 4A). In every matched pair, the PC version showed a larger fasting effect. For example, PC Hannum (β = −2.03 years, FDR p = 0.012) versus Hannum (β = −1.37 years, FDR p = 0.088); PC PhenoAge (β = −1.67 years, FDR p = 0.074) versus PhenoAge (β = −1.19 years, FDR p = 0.467); PC Horvath (β = −1.22 years, FDR p = 0.012) versus Horvath (β = +0.21 years, FDR p = 0.775).

**Figure 4.**
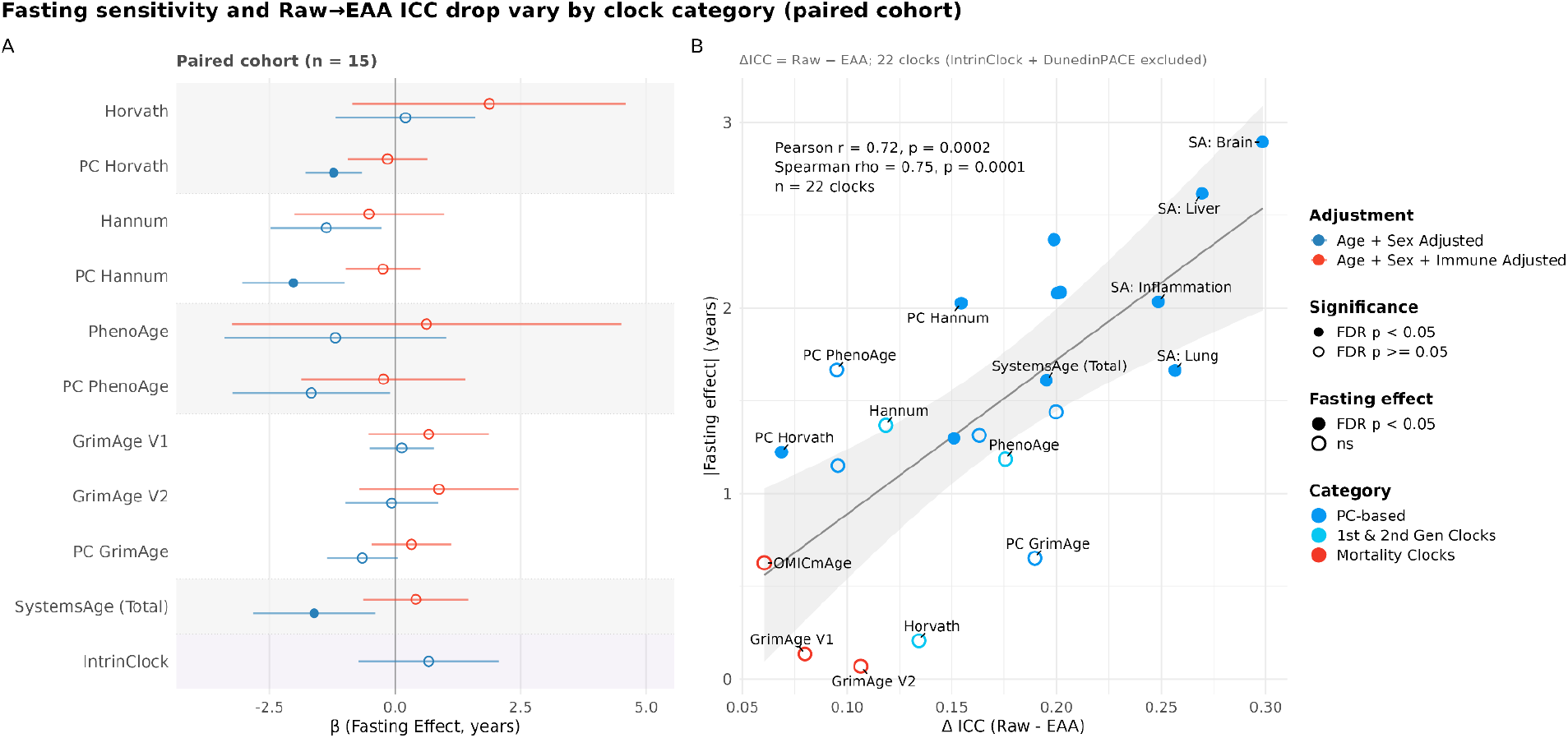
Fasting Effects in Matched PC vs Classical Clock Pairs and the Relationship Between ΔICC and Fasting Effect Size — Paired Cohort (n = 15). (A) Forest plot of fasting effects (β coefficients, years; 95% CI) for matched PC and classical clock pairs (Horvath / PC Horvath, Hannum / PC Hannum, PhenoAge / PC PhenoAge, GrimAge V1 / GrimAge V2 / PC GrimAge), with SystemsAge Total and IntrinClock shown for reference. Blue = EAA model (age and sex adjusted); red = IAA model (additionally adjusted for immune cell types). Filled symbols indicate FDR p < 0.05; open symbols indicate FDR p ≥ 0.05. Reference group: Non-Fasted. In every matched pair, the PC version showed a larger absolute fasting effect than its classical counterpart. (B) Scatter plot of the Raw → EAA ICC change (ΔICC) against the absolute fasting effect size (|β_EAA|, years) across 22 clocks (IntrinClock and DunedinPACE excluded). Each point is one clock, coloured by training category (PC-based, 1st and 2nd generation classical, Mortality). Symbol fill indicates whether the clock had an FDR-significant fasting effect at EAA. Shaded band = linear fit with 95% CI.

The PC transformation projects each classical clock’s CpG sites into a lower-dimensional space that captures correlated methylation variation more efficiently than the original clock coefficients ^13^. Immune cell composition is one of the largest sources of correlated CpG variance in whole blood, so the consistent PC > classical pattern is consistent with the PC transformation concentrating immune-correlated signal alongside the age signal each clock was originally trained on.

We then compared the fasting effect to the Raw-to-EAA ICC change (ΔICC) for 22 of the clocks (IntrinClock and DunedinPACE excluded). The ΔICC was correlated with the absolute size of the fasting effect (Pearson r = 0.72, 95% CI [0.43, 0.88], p = 1.6 × 10^−4^; Spearman ρ = 0.75, p < 10^−4^; Figure 4B). All three mortality clocks (OMICmAge ΔICC = 0.060, GrimAge V1 = 0.080, GrimAge V2 = 0.106) sat among the six smallest drops and had no FDR-significant fasting effect, and seven of the eight largest drops were SystemsAge organ clocks, with SA: Brain (ΔICC = 0.298), SA: Liver (0.269), SA: Lung (0.257), and SA: Inflammation (0.249) at the extreme (the eighth, SA: Immune, narrowly missed FDR at EAA: β = −1.44, FDR p = 0.064). Restricted to the 10 non-SystemsAge clocks, the correlation was weak (Pearson r = 0.18, p = 0.62).

Within the matched PC-vs-classical pairs themselves, ΔICC did not consistently mirror the fasting effect. PC Horvath (β = −1.22 years, ΔICC = 0.069) showed a smaller drop than Horvath (β = +0.21 years, ΔICC = 0.134) and PC PhenoAge (β = −1.67 years, ΔICC = 0.095) a smaller drop than PhenoAge (β = −1.19 years, ΔICC = 0.176) despite a larger fasting effect in both instances. Among the first and second generation matched pairs, only PC Hannum showed a larger drop than its classical counterpart. Why PC clocks may display smaller ΔICC despite larger fasting effects and how PC-driven variance compression contributes is discussed in the section 4.2.

### 3.4 Immune cell redistribution mediates the fasting signal

We have established that PC-clocks show the largest fasting effects relative to an acute refeeding and these effects are eliminated by immune cell adjustment. We have also noted that PC transformation amplifies each of the first and second generation classical clock’s fasting effects. Both lines of evidence point to immune cell changes as the mediator. We tested this directly in three ways: a) measuring the individual immune cell shifts that occur between fasted and fed states, b) quantifying each clock’s dependence on immune cell composition, and c) testing whether each clock’s immune context is correlated with the magnitude of the attenuation of the fasting effect across clocks.

In the paired cohort, fasting significantly increased CD4+ T naive cells (+54.8%, p = 0.004, FDR p = 0.048) and T regulatory cells (+42.7%, p = 0.008, FDR p = 0.048; within-person trajectories shown in Figure 5B), with trends toward decreased neutrophils (−7.2%, p = 0.089) and other shifts consistent with lymphocyte redistribution during fasting (Table 4, Figure 5A). The cross-sectional cohort confirmed this pattern with greater statistical power. Neutrophils showed the largest effect (−3.1%, FDR p < 0.001), 6 of the 12 cell types reached FDR significance (CD4+ T naive, CD8+ T naive, basophils, eosinophils, NK, neutrophils), and directional concordance between cohorts held for 9 of 12 cell types.

**Table 4.**
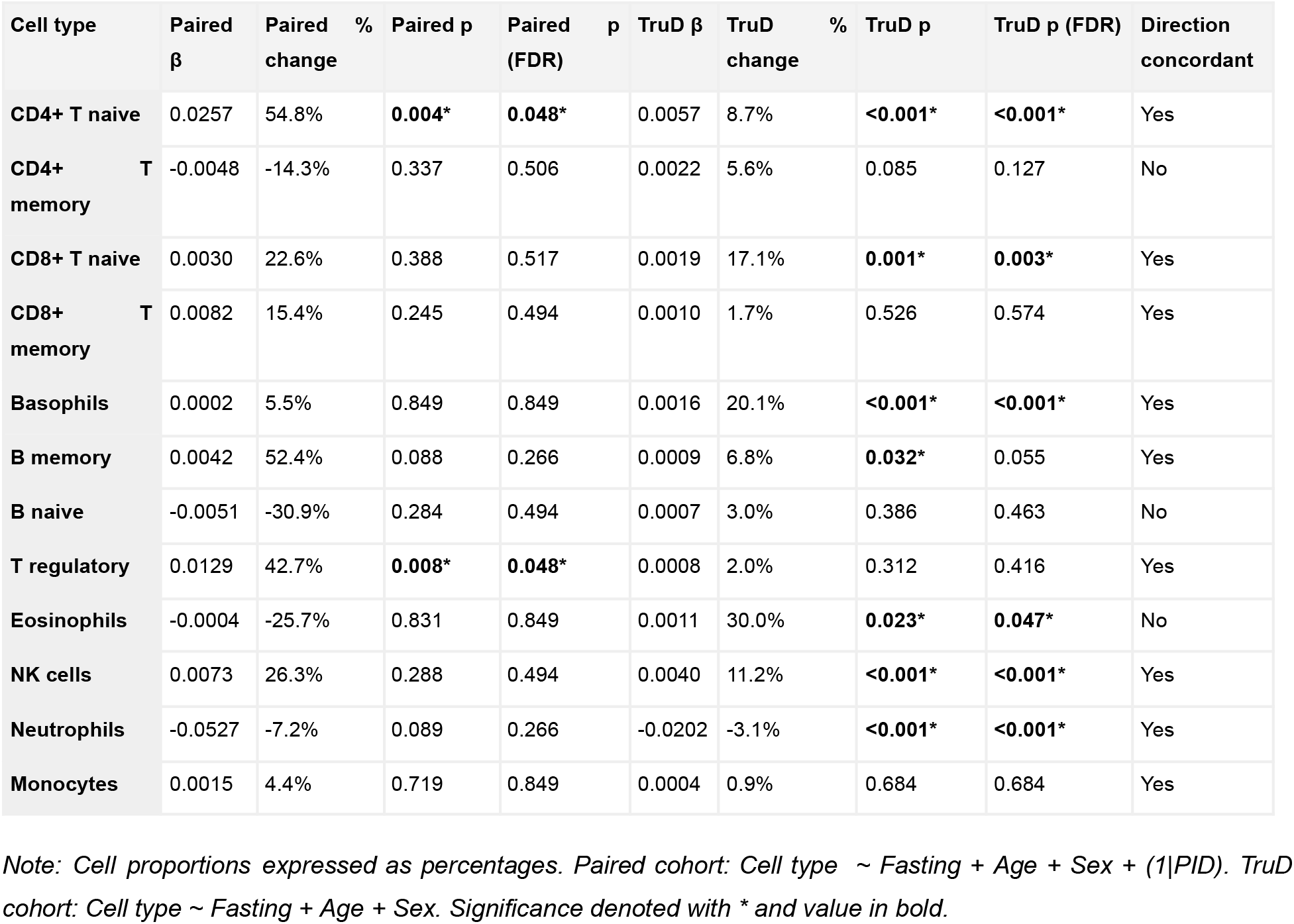
Immune Cell Composition by Fasting Status — Both Cohorts.

**Figure 5.**
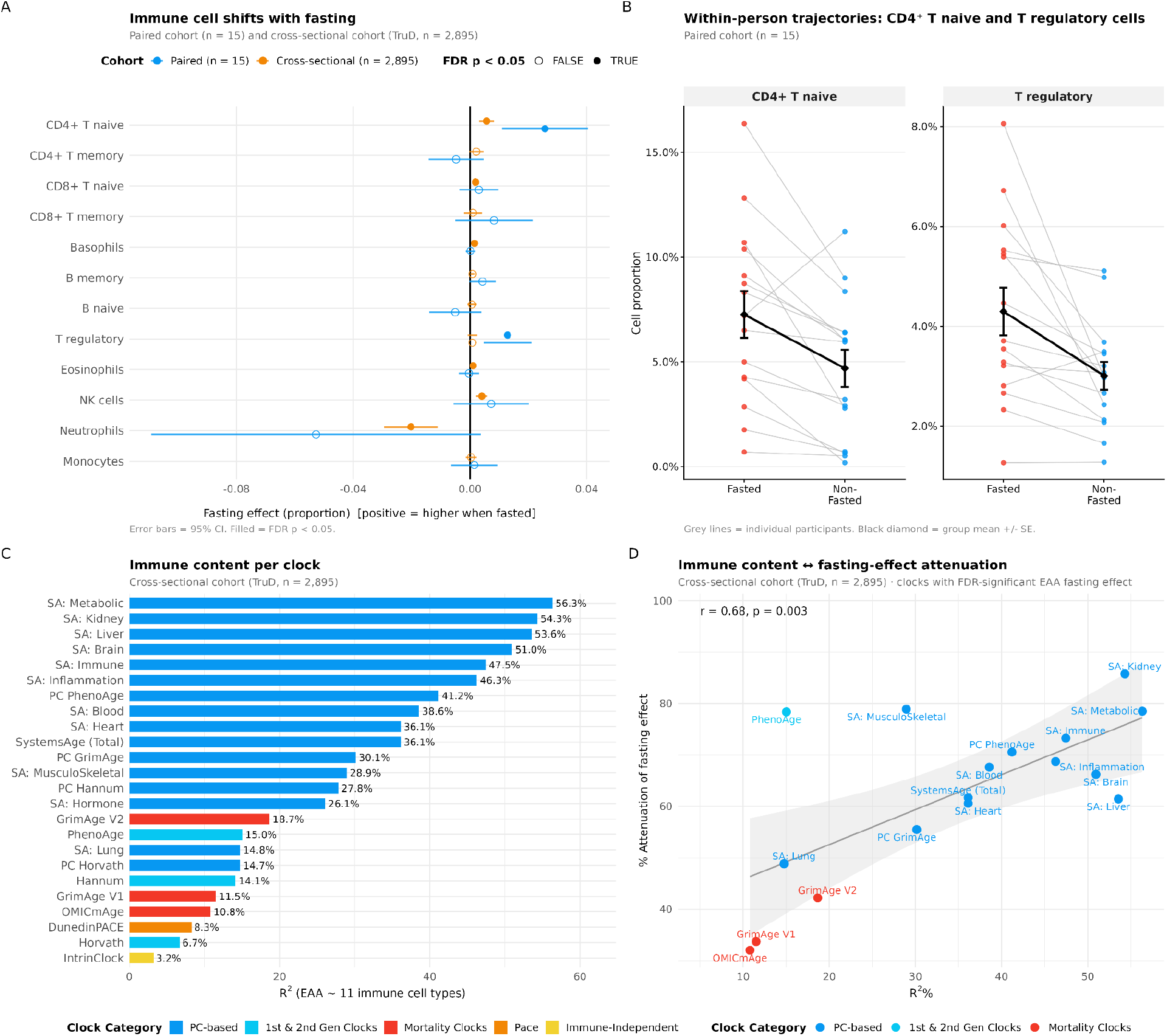
Immune Cell Composition Changes and Fasting Sensitivity. (A) Forest plot of fasting effects (β coefficients; 95% CI) on 12 immune cell proportions in the paired cohort (blue; linear mixed-effects model with random participant intercept) and cross-sectional cohort (orange; linear regression). Reference group: Non-Fasted; positive β = higher proportion when fasted. Filled symbols = FDR p < 0.05; open symbols = FDR p ≥ 0.05. (B) Within-person trajectories of CD4+ T naive cells and T regulatory cells across fasting states in the paired cohort, illustrating the two cell types with FDR-significant paired-cohort effects. Grey lines connect repeated measurements within the same participant; black diamond = group mean ± SE. (C) Fraction of age- and sex-residualised clock variance (R^2^) explained by 11 immune cell types, computed in the cross-sectional cohort (n = 2,895). Clocks are ordered by R^2^ and coloured by training category. PC-based and SystemsAge clocks show the highest immune content; mortality-trained clocks show intermediate content; IntrinClock shows the lowest. (D) Immune R^2^ correlates to fasting sensitivity: the fraction of EAA variance explained by immune cells correlates with the percent attenuation of the fasting effect by immune adjustment in the cross-sectional cohort (Pearson r = 0.68, p = 0.003). Each point is one clock with an FDR-significant cross-sectional EAA fasting effect; colours indicate clock category. Shaded band = linear fit with 95% CI.

To quantify each clock’s dependence on immune cell composition directly, we computed incremental R^2^ attributable to immune composition (Supplementary Table S2, Figure 5C). PC-based clocks showed the highest immune content. SystemsAge sub-clocks ranged from 14.8% (Lung) - 56.3% (Metabolic), and the PC transformations of first and second generation classical clocks ranged from 14.7% (PC Horvath) to 41.2% (PC PhenoAge). Their classical counterparts showed substantially lower values (Horvath: 6.7%, Hannum: 14.1%, PhenoAge: 15.0%). The immune R^2^ values reflect this pattern quantitatively. PC PhenoAge has approximately 2.75x the immune content of PhenoAge, and PC Horvath approximately 2x that of Horvath. Mortality clocks occupied an intermediate position (GrimAge V1 11.5%, GrimAge V2 18.7%, OMICmAge 10.8%), while IntrinClock had the lowest immune content at 3.2% as expected.

Within the cross-sectional cohort, immune R^2^ was correlated with the degree to which immune adjustment attenuated the fasting effect across clocks (Pearson r = 0.68, p = 0.003, n = 17 clocks with FDR-significant EAA fasting effects; DunedinPACE was excluded because % attenuation is not comparable for a pace-of-aging clock; Figure 5D). Because both quantities are computed in the same cohort, the relationship is a within-cohort association consistent with immune cell composition being a source of the observed clock shifts; it does not independently test paired-cohort effects, and the correlational design cannot establish causation. Ten of these 17 clocks are SystemsAge organ sub-clocks (SystemsAge: Hormone excluded as it did not reach EAA significance), which are highly intercorrelated (mean Spearman ρ ≈ 0.92 across sub-clock pairs). Treating SystemsAge as a single observation reduces the scatter to n = 7 clock categories, where the correlation is directionally consistent but no longer statistically significant (r = 0.57, p = 0.18). The association is therefore best interpreted as a pattern across clock categories rather than a robust per-clock dose-response.

Cross-sectional immune adjustment produced only partial attenuation of mortality-clock effects (32–42%, versus ≥74% for the immune-mediated clocks in the paired cohort), but no clock retained an FDR-significant fasting effect after adjustment (all FDR p ≥ 0.07). This hints at self-reported cross-sectional fasting status potentially capturing biology beyond acute immune redistribution, likely lifestyle and metabolic confounding between habitual fasters and non-fasters, that the within-person paired design isolates out.

IntrinClock provides the negative control for this mechanism. Its fasting association was null in both cohorts (Table 1, Table 2, Figure 2 A,B), consistent with its immune R^2^ of 3.2%. If fasting effects on clock outputs were driven by a mechanism other than immune cell redistribution, such as metabolic changes affecting methylation directly, IntrinClock should also be affected. Its null result supports immune cell composition as the primary mediator of fasting effects on clock outputs.

## 4. Discussion

### 4.1 Clock construction as an organizing principle in fasting response

A key observation is that a clock’s response to fasting varies systematically with how the clock was built. Within the cross-sectional cohort, a clock’s immune R^2^, the fraction of its age- and sex-residualised variance explained by immune cell composition, is correlated with the degree to which immune adjustment attenuates its fasting effect. The correlational design cannot establish causation, and both quantities are computed in the same cohort, so the relationship is a within-cohort association rather than an independent test. With that caveat, the observed pattern across clock categories is consistent with the following reading.

PC-based clocks (immune R^2^ 15–56%) are the most sensitive, because the PC transformation projects CpG methylation into a space that efficiently captures cell-composition variance alongside aging biology, thereby concentrating the immune signal. SystemsAge, built natively from principal components, is the most extreme example. First- and second-generation classical clocks (immune R^2^ 7–15%) show intermediate sensitivity, but their PC transformation roughly triples the immune dependence of the same underlying clock, as developed in detail below.

Mortality-trained clocks (immune R^2^ ~11–19%) are the least sensitive to fasting in the paired cohort, because their construction routes through DNAm surrogates of plasma proteins (GrimAge) or multi-omic features (OMICmAge) whose between-person variation reflects chronic health rather than acute immune shifts. PC GrimAge is an informative special case, as its immune R^2^ is roughly triple that of original GrimAge, but its paired-cohort fasting effect did not reach significance at n = 15 and its IAA ICC barely shifted from EAA. One reading is that the mortality-training target contributes a non-immune component that moderates the fasting response on top of the PC architecture, but with n = 15 we cannot rule out limited power as the explanation. Sehgal et al. identified PC GrimAge as the only clock achieving good biological reliability (ICC > 0.75) in their pooled analysis ^9^, consistent with our observations, though direct replication in a larger paired design would be needed to confirm the proposed mechanism.

DunedinPACE was resistant to fasting in the paired cohort despite intermediate immune content. Unlike age-prediction clocks, it measures the pace of biological aging from longitudinal within-person change, making it inherently less sensitive to cross-sectional composition shifts. IntrinClock has the lowest immune content by design and serves as a negative control for the entire analysis.

This framework has a practical implication that the field has not fully appreciated. The question as to whether epigenetic clocks are reliable has no single answer. It depends on which clock, which perturbation, and what aspect of biology the clock was constructed to capture. We agree with Sehgal et al. ^9^ that a single meal or acute stressor should not truly age or de-age an individual by several years. However, this observation supports our argument rather than undermining it, which is that the clocks reporting such shifts are detecting real changes in immune cell composition, not changes in biological age. The appropriate correction for affected clocks is immune cell adjustment, which recovers the ICC by removing perturbation-driven variance (e.g., SystemsAge). For mortality-trained clocks, no such correction is needed in the acute context, as their architecture routes through DNAm surrogates of plasma proteins (GrimAge), or of proteins, metabolites, and clinical measures (OMICmAge) - features whose variance may reflect chronic biology rather than transient immune redistribution.

The clocks most sensitive to this effect employ principal component analysis at multiple stages of their construction. SystemsAge performs PCA on system-specific biomarkers, PCA on CpG methylation data, elastic net regression from methylation PCs to system biomarker PCs, and a final PCA on the resulting system scores before mortality prediction ^6^. At each step, PCA concentrates correlated variation, and immune cell composition is among the largest sources of correlated CpG variation in whole blood. The PC transformation of first- and second-generation classical clocks operates on the same principle, projecting each clock’s CpG sites into a lower-dimensional space that captures shared methylation patterns ^13^. Our data quantify the consequence. The PC transformation approximately doubles or triples the fraction of clock variance attributable to immune composition, a consequence of the PC architecture. A clock that registers fasting-driven immune changes is behaving as a sensitive instrument, so the relevant question is not whether such a clock is reliable in the abstract but whether the sampling context has been accounted for.

This reframing connects to a broader principle from clinical laboratory medicine, whereby useful biomarkers routinely tolerate meaningful biological variation when the variation is understood. HDL cholesterol, for example, is assessed against a total allowable error (TAE) threshold of roughly 20% and remains clinically valuable precisely because its fluctuations are interpreted within physiological and clinical context. Although ICC and TAE are not directly comparable metrics, the parallel is that a clock detecting real physiological change is not “unreliable” in a way that disqualifies it from use, rather, it is sensitive and requires the sampling context to be understood and controlled. We develop the practical consequences of this view.

### 4.2 Interpreting ICC for biological reliability

ICC is an excellent metric for technical reliability, but its utility for biological reliability requires accounting for cohort structure, perturbation type and its mechanism, and the adjustment level (e.g. raw output, age-residualised, etc). Since ICC is between-person variance divided by total variance (between + within), it is sensitive both to within-person noise and to the spread of between-person variance in the cohort. We unpack four scenarios in which ICC alone falls short of capturing biological reliability.

First, high Raw ICCs can coexist with systematic group-level shifts. A fasting shift of 0.5 - 3 years in our paired cohort is small relative to the age-dominated between-person spread in Raw output (ages 24–52, SD = 8.4 years), so Raw ICCs remain ≥ 0.96 even as effect models detect structured directional biology in the same clocks. ICC and group-level effect estimates therefore answer different questions, and ICC alone does not characterise a clock’s behaviour under a specific perturbation.

Second, ICCs must be evaluated in a cohort- and perturbation-specific manner. Different perturbations operate through different pathways, affect different cell types, and elicit different clock responses. A fasting analysis altering immune composition will not produce comparable ICCs to a stress or environmental exposure study, and biological sensitivity to one perturbation can be mistaken for measurement instability when the source of within-person variance is not characterised.

Third, the per-clock Raw-to-EAA ICC drop is confounded by PC architecture. The ICC drop reflects two competing forces. When fasting shifts a clock, within-person variance rises and ICC falls a real biological signal. But the PC transformation independently compresses within-person variance, dampening the ICC drop even when a biological effect is present ^13^. PC Horvath illustrates this directly: its EAA ICC was higher than the matched classical Horvath clock, despite a larger fasting effect, because PC compresses within-person variance more than between-person variance (Table 3). A small Raw-to-EAA drop therefore does not unambiguously indicate fasting insensitivity, as it could equally reflect noise compression masking a real signal. The category-level pattern is, however, interpretable, whereby immune-sensitive clocks show both the largest group-level effects and the largest Raw-to-EAA drops, while mortality-trained clocks show neither.

Fourth, ICC declines structurally with each adjustment level. Each successive adjustment, from Raw to EAA to IAA, removes structured between-person variance from the denominator, producing a mathematical expectation that ICC will decline with each step. From Raw to EAA, the age-dominated between-person spread shrinks and fasting-driven within-person variance has room to register, which is why EAA ICCs decline to a heterogeneous good-to-excellent range. From EAA to IAA, fitting 11 immune cell proportions can raise or drop ICC depending on whether it is removing or adding within-person variance. In our technical replicate analysis which serves as a useful technical noise anchor, we show IAA ICCs decline by a median of 0.73 from EAA even in the absence of any biology between paired measurements.

Through this lens, our study extends Sehgal et al.’s biological reliability assessment in three respects ^9^. First, rather than pooling ICCs across mechanistically distinct perturbations and array platforms, which may conflate sources of variance, we report on a single, well-characterised perturbation, in which the sources of within-person variance can be isolated and interpreted. Second, their reported biological reliability ICCs are age-residualised (EAA). As mentioned above, age residualisation removes the between-person age variance, so EAA ICCs are expected to be substantially lower than raw-clock ICCs on the same data, a structural consequence of the adjustment. We decompose reliability across successive adjustment levels to gauge the contribution of an adjustment to the reported value. Third, the paper identified immune cell dynamics, stress pathways, and metabolic fluctuations as candidate sources of biological variability. We test this directly for fasting, and demonstrate that the relevant mechanism (e.g. immune cell redistribution) can be accounted for analytically rather than read as a reliability failure. More broadly, judging a clock unsuitable solely because its ICC falls below a fixed threshold risks mistaking sensitivity for noise.

### 4.3 Immune adjustment is not a default correction

A critical insight from our analysis is that immune cell adjustment should not be treated as a uniform correction step applied to all clocks. Whether immune-mediated clock changes represent “signal” or “confound” depends entirely on the analytical context. In studies examining the aging consequences of immune cell redistribution, such as infection, vaccination, immunotherapy or even chronic disease, the immune component may be the signal of interest and should not be adjusted away. In studies asking whether an intervention affects aging biology beyond immune changes, immune adjustment isolates the residual and is appropriate.

Immune adjustment is also clock-specific in its effect. To interpret ICC under immune adjustment when the perturbation itself changes immune composition, we have to break ICC into its between-person and within-person components and ask which is changing with each successive adjustment, why, and what a high or low IAA ICC tells us about the utility of the ICC after adjustment, or about whether the adjustment was appropriate in the first place. In the fasting pairs, this adjustment (which fits 11 immune cell types to account for all 12) has two competing effects on within-person variance: a) it can remove perturbation-driven variance (raising ICC), or it can add estimation noise by fitting 11 heterogeneous immune profiles to relatively stable residuals (lowering ICC). Which effect dominates determines the pattern, as reported in section 3.2.

When ICC recovers after immune adjustment as within-person variance is removed, (SystemsAge Total), the immune adjustment in this scenario may be appropriate if the goal is to ask whether fasting-related changes exist beyond immune cell composition. When the residuals are already stable across fasting states and adjustment adds within-person variance by fitting noisy immune profiles to them, ICC collapses (OMICmAge). Here, immune adjustment may be inappropriate, since it is injecting estimation noise without removing meaningful perturbation signals. When a clock measures purely immune biology (SystemsAge: Immune), IAA ICC collapses entirely. In this case adjustment serves as a positive control but IAA ICC is uninterpretable as a stability metric, since no non-immune signal remains to evaluate. Each pattern is a diagnostic of clock composition.

The mortality clocks fit this gradient. GrimAge V1 and V2 predict mortality through DNAm surrogates of plasma proteins, some of which (e.g. CRP in GrimAge V2) are themselves immune-correlated; their IAA ICCs decline moderately, reflecting the partial cost of fitting immune-related features into the residual. OMICmAge collapses more dramatically, and from both sides. In addition to the within-person noise described above, immune adjustment removes 78% of its between-person variance, compared with ~48% for GrimAge V1, suggesting that its multi-omic construction embeds immune-correlated signals more deeply into its between-person structure. PC GrimAge, by contrast, shows almost no ICC change (0.791 to 0.754) because its non-immune mortality biology survives adjustment as a stable residual.

Immune adjustment and ICC are difficult to interpret jointly when the perturbation itself changes immune profiles. Variance decomposition becomes necessary in this setting, as it is the only way to tell whether a low or high IAA ICC reflects clock stability, adjustment overfitting, or a clock that was measuring immune biology all along.

### 4.4 Practical Implications

#### For Intervention Trials

Our results have direct implications for the design and interpretation of aging intervention trials. Fasting status must be standardized at all blood collection timepoints. A participant who provides a fasted sample at baseline and a fed sample at follow-up could show a spurious epigenetic age increase of 1 – 3 years, potentially masking a genuine intervention effect. Conversely, the reverse pattern could produce a false-positive result. We recommend that all aging studies specify and report fasting status as part of their sample collection protocol.

When fasting status cannot be controlled, immune-adjusted (IAA) models should be used as a sensitivity analysis. However, researchers should be cautious about defaulting to IAA for mortality clocks, particularly OMICmAge, where immune adjustment substantially reduces the variance rather than removing the confounding noise. A practical recommendation is to present both EAA and IAA results and discuss the implications of any discrepancies.

#### For Study Design

In large cohort studies where fasting status may vary, our findings suggest that fasting status should be explicitly considered in study design or analysis. This can involve standardizing collection conditions, modeling fasting as a covariate, or examining clock values with and without immune cell adjustment, depending on the biological question. Given the likely mechanism that fasting is influencing DNA methylation through immune redistribution, simultaneous adjustment for both fasting and immune cell composition may obscure the biological pathway of interest. In such context, models should be structured to reflect the assumed causal relationships rather than applied automatically. When fasting status is unavailable, time of collection may serve as a proxy (morning samples are more likely to be fasted). The

TruDiagnostic validation demonstrates that even self-reported fasting status, despite its imprecision, captures meaningful variation in clock outputs. Importantly, immune cell adjustment should not be treated as a uniform correction step, but as an informed strategy that alters the biological question being asked.

#### For Model Development

Any predictive model should consider the degree to which their algorithms are sensitive to immune cell composition as a design dimension. Clocks trained on mortality or pace-of-aging targets, such as GrimAge V1/V2, OMICmAge, and DunedinPACE, were robust to acute refeeding following an overnight fast in our paired cohort, and may be preferable when fasting status cannot be controlled. Clocks whose ICC remains stable under immune adjustment, such as PC GrimAge, may also be preferable in this setting. By contrast, SystemsAge and other PC-based clocks are highly sensitive to immune redistribution, which requires controlled sampling. Future model development could explicitly optimize for robustness to immune cell variation (as IntrinClock does), balanced against the biological relevance of immune-related aging signals.

### 4.5 Relationship to Prior Work

Our findings build on and extend several prior observations. Zhang et al. ^12^ demonstrated in a cross-sectional analysis that cell type composition explains a large proportion of EAA variation, establishing the theoretical foundation for our work. Our fasting study provides a dynamic, within-person experimental confirmation of this relationship: when immune cells change acutely due to fasting, clocks respond accordingly. Guo et al. ^21^ showed that innate immune cell subtypes, including neutrophils and NK cells, correlate with epigenetic clock outputs in population studies. Our data show that these same cell types are among the most responsive to fasting (neutrophils p = 1.6 × 10^−5^ and NK cells p = 4.4 × 10^−5^ in TruD), providing a mechanistic link between acute immune perturbation and clock behavior.

With respect to Sehgal et al. ^9^ specifically, our study builds on their analysis by adding a mechanistic interpretation of biological reliability using ICC for a single perturbation. Moreover, we show how clock construction shapes the within-person variance, and how variance decomposition makes that variance interpretable rather than reading a reduced ICC as instability.

### 4.6 Limitations

Several limitations should be acknowledged. First, our paired cohort is small (N = 15), limiting statistical power for detecting smaller effects. While the within-person design partially compensates by eliminating between-person variability, some true fasting effects may have been missed, particularly for immune cell types that showed large but non-significant trends. Second, the TruDiagnostic validation uses a cross-sectional design with self-reported fasting status at a single timepoint, which introduces measurement error, cannot distinguish overnight fasting from habitual fasting practice, and prevents within-person ICC estimation. Mortality-trained clocks reached EAA significance at population scale in the cross-sectional cohort but did not survive immune adjustment, and cross-sectional immune attenuation was consistently lower than paired attenuation; we therefore interpret the cross-sectional cohort as confirming the training-category sensitivity gradient rather than as evidence that fasting status causally moves mortality-clock outputs at the population level. Residual lifestyle, metabolic, or collection-timing differences between self-reported fasters and non-fasters likely contribute. Third, immune cell proportions were estimated by DNA methylation deconvolution rather than measured directly (e.g., flow cytometry or a complete blood count); a future cohort that repeats the fasting and refeeding design with directly measured cell counts is needed to confirm that the deconvolution-estimated shifts reflect true changes in circulating immune composition. Relatedly, because immune composition and metabolic state are coupled during fasting and refeeding, immune adjustment removes shared collinear variance and should be read as decomposition rather than as proof of immune causation; the immune-specificity of the effect instead rests on the negative-control and dose-response results above. Fourth, we examined only overnight fasting (12+ hours) followed by acute refeeding; prolonged fasting, intermittent fasting protocols, and dietary composition effects were not evaluated. Fifth, we did not perform formal causal mediation analysis; our change-in-effect approach demonstrates attenuation but does not establish formal mediation in the causal inference framework.

### 4.7 Future Directions

Several extensions of this work would be valuable. Time-course studies with serial blood draws at multiple intervals following a meal would characterize the kinetics of immune cell and clock changes, informing practical fasting windows for sample collection. Investigation of dietary composition effects (e.g., high-fat vs. high-carbohydrate meals) could reveal whether specific macronutrients differentially influence immune cell proportions and clock outputs. Individual differences in fasting sensitivity, potentially related to age, sex, metabolic health, or genetic variation, could inform personalized sampling protocols. Finally, extending this framework to other transient physiological perturbations, such as exercise, stress, jet lag or sleep deprivation, would more comprehensively characterize the pre-analytical factors that influence epigenetic clock measurements.

## 5. Conclusion

Epigenetic clocks demonstrated excellent within-person stability across fasting states, with Raw ICCs of 0.93 – 0.98 across all clocks except DunedinPACE (Raw ICC = 0.81). The systematic mean shifts observed between fasted and non-fasted samples, ranging from 0.5 to 3.0 years across immune-sensitive clocks, are not evidence of clock instability, but reflect appropriate sensitivity to genuine, transient immune cell composition changes. Adjusting for methylation-estimated immune cell types attenuated 74–100% of FDR-significant fasting effects in the paired cohort, with most clocks above 80% attenuation, identifying immune cell composition as the primary mechanistic pathway through which fasting influences epigenetic age estimates. No clock retained an FDR-significant fasting effect at the IAA level in either the paired (n = 15) or the cross-sectional (n = 2,895) cohort.

These findings replicated in the cross-sectional cohort of 2,895 individuals, confirming the generalisability of the immune-mediation pattern at population scale. Mortality-trained clocks (GrimAge V1/V2, OMICmAge) and DunedinPACE were robust to fasting in the paired design, consistent with construction targets (i.e. long-term mortality risk and pace-of-aging) that do not move under acute immune redistribution. PC GrimAge was a notable case among PC-based clocks, whereby it showed the smallest paired-cohort fasting effect of the PC clocks and the only IAA ICC that was essentially unchanged from EAA, consistent with non-immune mortality biology surviving immune adjustment.

Our study calls for three practical changes: (1) standardisation of fasting status in all epigenetic aging studies; (2) reporting of both age-adjusted (EAA) and immune-adjusted (IAA) clock values as complementary measures, with attention to clock-specific behaviour under immune adjustment, and (3) reinterpretation of ICC-based reliability assessments for biological reliability as cohort- and perturbation-specific quantities, with explicit benchmarking against technical reliability. Rather than undermining the utility of epigenetic clocks, understanding the immune cell contribution clarifies what they measure and how to use them appropriately in research and clinical contexts.

