## Supplementary Tables and Figures for "Fasting Status and Epigenetic Clock Stability: Implications for Aging Research" (from another version of this preprint: version 1, 2026-06-06)

##### Supplementary Table 1. Intraclass Correlation Coefficients (ICC) Across Adjustment Levels — Technical Replicates EPICv2custom (n = 4 x 24 replicates)

| Clock | Raw ICC (95% CI) | EAA ICC (95% CI) | IAA ICC (95% CI) | ΔRaw→EAA |
|---|---|---|---|---|
| **PCHorvath** | 0.979 (0.936–0.998) | 0.928 (0.797–0.994) | 0.273 (0.082–0.851) | 0.051 |
| **PCHannum** | 0.988 (0.961–0.999) | 0.968 (0.902–0.998) | 0.342 (0.118–0.886) | 0.020 |
| **PCPhenoAge** | 0.977 (0.928–0.998) | 0.856 (0.642–0.988) | 0.147 (0.027–0.742) | 0.121 |
| **PCGrimAge** | 0.998 (0.992–1.000) | 0.870 (0.670–0.989) | 0.078 (0.001–0.623) | 0.128 |
| **Horvath** | 0.866 (0.662–0.989) | 0.537 (0.247–0.944) | 0.026 (-0.016–0.448) | 0.329 |
| **Hannum** | 0.943 (0.836–0.996) | 0.668 (0.371–0.966) | 0.111 (0.010–0.693) | 0.275 |
| **PhenoAge** | 0.782 (0.517–0.981) | 0.404 (0.153–0.909) | 0.006 (-0.025–0.360) | 0.378 |
| **GrimAge V1** | 0.949 (0.851–0.996) | 0.750 (0.471–0.977) | 0.142 (0.024–0.736) | 0.199 |
| **GrimAge V2** | 0.940 (0.828–0.995) | 0.783 (0.519–0.981) | 0.232 (0.062–0.825) | 0.157 |
| **OMICmAge** | 0.974 (0.918–0.998) | 0.409 (0.158–0.910) | 0.013 (-0.021–0.392) | 0.565 |
| **SystemsAge** | 0.990 (0.967–0.999) | 0.937 (0.820–0.995) | 0.206 (0.049–0.806) | 0.053 |
| **SystemsAge Blood** | 0.983 (0.946–0.999) | 0.942 (0.831–0.996) | 0.201 (0.047–0.801) | 0.041 |
| **SystemsAge Brain** | 0.962 (0.887–0.997) | 0.805 (0.552–0.983) | 0.065 (-0.008–0.601) | 0.157 |
| **SystemsAge Inflammation** | 0.990 (0.967–0.999) | 0.909 (0.752–0.993) | 0.000 (-0.030–0.351) | 0.081 |
| **SystemsAge Heart** | 0.990 (0.968–0.999) | 0.910 (0.754–0.993) | 0.240 (0.066–0.831) | 0.080 |
| **SystemsAge Hormone** | 0.974 (0.921–0.998) | 0.902 (0.737–0.992) | 0.000 (-0.030–0.351) | 0.072 |
| **SystemsAge Immune** | 0.977 (0.929–0.998) | 0.934 (0.813–0.995) | 0.000 (-0.029–0.345) | 0.043 |
| **SystemsAge Kidney** | 0.991 (0.972–0.999) | 0.936 (0.817–0.995) | 0.000 (-0.027–0.328) | 0.055 |
| **SystemsAge Liver** | 0.966 (0.896–0.997) | 0.873 (0.674–0.990) | 0.051 (-0.013–0.564) | 0.093 |
| **SystemsAge Metabolic** | 0.984 (0.949–0.999) | 0.945 (0.840–0.996) | 0.016 (-0.023–0.428) | 0.039 |

| | | | | |
|---|---|---|---|---|
| **SystemsAge Lung** | 0.978 (0.933–0.998) | 0.858 (0.646–0.988) | 0.004 (-0.024–0.341) | 0.120 |
| **SystemsAge MusculoSkeletal** | 0.976 (0.924–0.998) | 0.868 (0.665–0.989) | 0.049 (-0.010–0.546) | 0.108 |
| **DunedinPACE** | — | 0.557 (0.263–0.948) | 0.109 (0.011–0.688) | — |
| **IntrinClock** | 0.912 (0.760–0.993) | 0.170 (0.036–0.769) | — | 0.742 |

*Note: ICC computed using two-way mixed-effects model (ICC 2,1). EAA = Epigenetic Age Acceleration; IAA = Immune-Adjusted Acceleration. DunedinPACE is a pace-of-aging measure and does not have an EAA variant.*

**Supplementary Table 2. Immune cell composition explains epigenetic age acceleration variance — EAA ~ 11 immune cell types (TruDiagnostic cross-sectional cohort, n = 2,895).**

| Clock | $R^2$ | Adjusted $R^2$ | $R^2$ (%) | F-test p |
|---|---|---|---|---|
| **PC Horvath** | 0.147 | 0.144 | 14.7% | <0.001 |
| **PC Hannum** | 0.278 | 0.275 | 27.8% | <0.001 |
| **PC PhenoAge** | 0.412 | 0.410 | 41.2% | <0.001 |
| **PC GrimAge** | 0.301 | 0.299 | 30.1% | <0.001 |
| **Horvath** | 0.067 | 0.064 | 6.7% | <0.001 |
| **Hannum** | 0.141 | 0.138 | 14.1% | <0.001 |
| **PhenoAge** | 0.150 | 0.147 | 15.0% | <0.001 |
| **GrimAge V1** | 0.115 | 0.112 | 11.5% | <0.001 |
| **GrimAge V2** | 0.187 | 0.184 | 18.7% | <0.001 |
| **OMICmAge** | 0.108 | 0.105 | 10.8% | <0.001 |
| **SystemsAge (Total)** | 0.361 | 0.359 | 36.1% | <0.001 |
| **SA: Blood** | 0.386 | 0.383 | 38.6% | <0.001 |
| **SA: Brain** | 0.510 | 0.508 | 51.0% | <0.001 |
| **SA: Inflammation** | 0.463 | 0.461 | 46.3% | <0.001 |
| **SA: Heart** | 0.361 | 0.359 | 36.1% | <0.001 |
| **SA: Hormone** | 0.261 | 0.258 | 26.1% | <0.001 |
| **SA: Immune** | 0.475 | 0.473 | 47.5% | <0.001 |
| **SA: Kidney** | 0.543 | 0.541 | 54.3% | <0.001 |
| **SA: Liver** | 0.536 | 0.534 | 53.6% | <0.001 |
| **SA: Metabolic** | 0.563 | 0.562 | 56.3% | <0.001 |

| | | | | |
|---|---|---|---|---|
| **SA: Lung** | 0.148 | 0.144 | 14.8% | <0.001 |
| **SA: MusculoSkeletal** | 0.289 | 0.287 | 28.9% | <0.001 |
| **DunedinPACE** | 0.108 | 0.104 | 10.8% | <0.001 |
| **IntrinClock** | 0.032 | 0.029 | 3.2% | <0.001 |

### Supplementary Figures

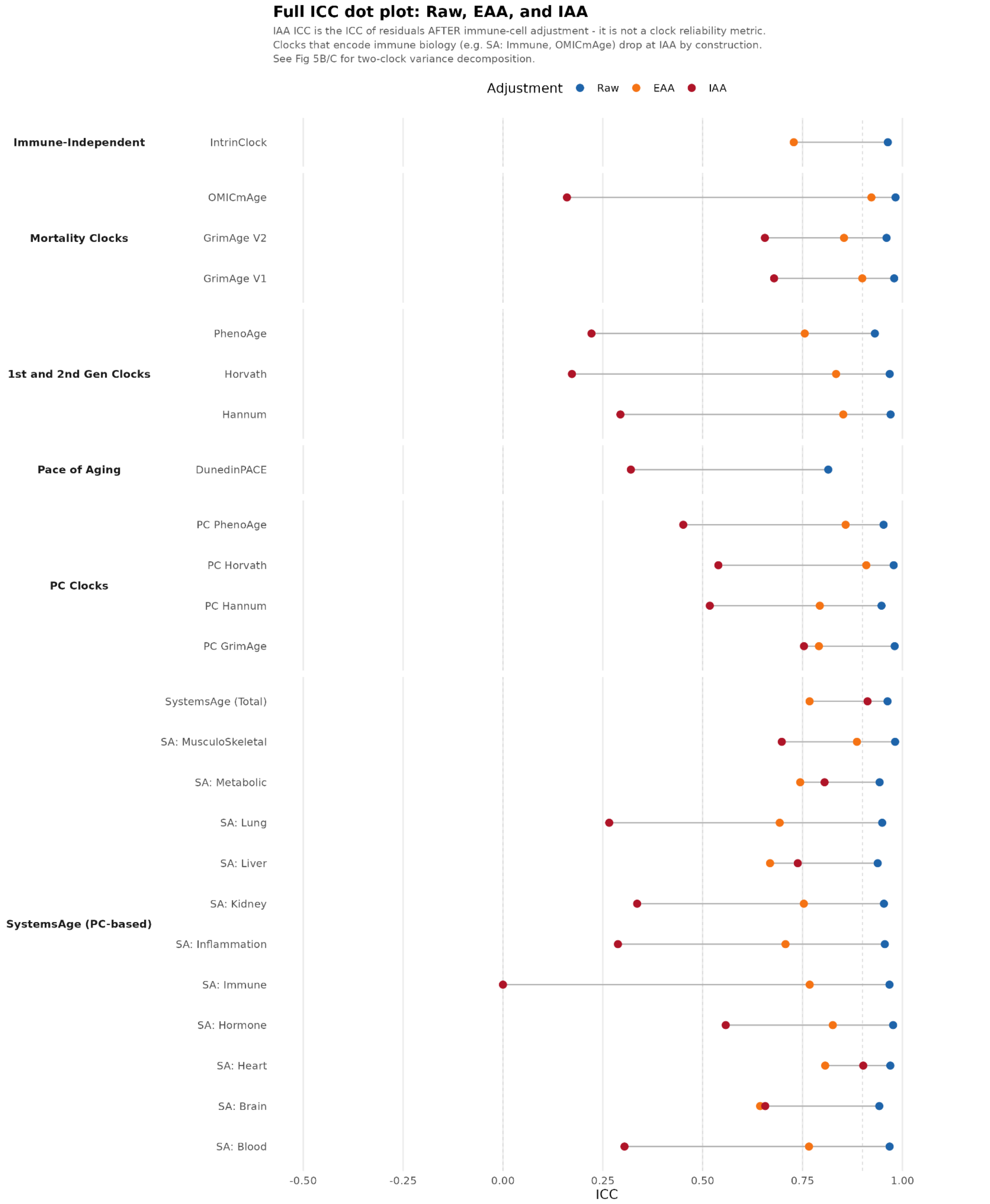


***Supplementary Figure S1. Test-Retest Reliability Across Three Adjustment Levels — Paired Cohort.*** *Intraclass correlation coefficients (point estimates) for all 24 clocks at three adjustment levels: Raw (blue), EAA (orange), and*

*IAA (red), grouped by training category. Raw ICC reflects the raw clock output; EAA removes age-driven between-person variance via residualisation on Age; IAA additionally removes immune-composition-driven between-person variance via residualisation on Age plus 11 immune cell types (Monocytes excluded to avoid collinearity). DunedinPACE is shown at Raw and IAA only (no chronological-age EAA variant); IntrinClock is shown at Raw and EAA only (modelled as a singleton immune-independent negative control).*
